# Binocular integrated visual field deficits are associated with changes in local network function in primary open-angle glaucoma: a resting-state fMRI study

**DOI:** 10.1101/2021.07.19.452985

**Authors:** Giorgia Demaria, Azzurra Invernizzi, Daniel Ombelet, Joana C. Carvalho, Remco J. Renken, Frans W. Cornelissen

## Abstract

Recent brain imaging studies have shown that the degenerative eye damage generally observed in the clinical setting, also extends intracranially. Both structural and functional brain changes have been observed in glaucoma participants, but we still lack an understanding of whether these changes also affect the integrity of cortical functional networks. This is relevant, as functional network integrity may affect the applicability of future treatments, as well as the options for rehabilitation or training. Here, we compare global and local functional connectivity between glaucoma and controls. Moreover, we study the relationship between functional connectivity and visual field (VF) loss.

For our study, 20 subjects with primary open-angle glaucoma (POAG) and 24 age-similar healthy participants were recruited to undergo a complete ophthalmic assessment followed by two resting-state (RS) (f)MRI scans. For each scan and for each group, the ROIs with EC values higher than the 95th percentile were considered the most central brain regions (“hubs”). Hubs for which we found a significant difference in EC in both scans between glaucoma and healthy were considered to provide evidence for network changes. In addition, for each participant, behavioural scores were derived based on the notion that a brain region’s hub function might relate to the: 1) sensitivity of the worse eye – indicating disease severity, 2) sensitivity of both eyes combined – with one eye potentially compensating for loss in the other, or 3) difference in eye sensitivity – requiring additional network interactions. By correlating each of these VF scores and the EC values, we assessed whether VF defects could be associated with centrality alterations in POAG. Our results show that no functional connectivity disruptions were found at the global brain level in POAG participants. This indicates that in glaucoma global brain network communication is preserved. Furthermore, a positive correlation was found between the EC value of the Lingual Gyrus, identified as a brain hub, and the behavioural score for the VF sensitivity of both eyes combined. The fact that reduced local network functioning is associated with reduced sensitivity of the binocular VF suggests the presence of local brain reorganisation that has a bearing on functional visual abilities.

## 1. Introduction

Glaucoma is a neurodegenerative ophthalmic disease, which is characterized, amongst others, by reduced retinal thickness and loss of visual field (VF) sensitivity. It is one of the leading causes of irreversible blindness, and in 2020 approximately 76 million people are affected by this disease, a number that is only estimated to increase in the coming years (Quigley & Broman, 2006; Tham et al., 2014). Thus, it is important to better understand the pathogenesis of glaucoma, in order to improve its diagnosis, prognosis and the possible treatment options, all of which will ultimately improve the quality of life of patients. While in the clinical setting the degenerative physical damage is generally observed in the eye and in the optic nerve, recent brain imaging studies have shown that this is not limited to these but extends intracranially along the visual pathways and into the visual cortex (Gupta & Yücel, 2007; Haykal et al., 2021; Hernowo, 2012; Nucci et al., 2013). It has been shown that structural and functional changes in the brain are involved in glaucoma. Structural changes were observed in several studies that compared glaucoma participants to healthy controls, showing altered white matter tracts and gray matter atrophy in brain areas involved in visual processing (Chen et al., 2013; Frezzotti et al., 2014; Giorgio et al., 2018; J. Wang, Li, Sabel, et al., 2016). Moreover, several functional magnetic resonance imaging (fMRI) studies have shown altered cortical activity in glaucoma patients (Gerente et al., 2015; Jiang et al., 2017; Qing et al., 2010; Y. Wang et al., 2020). It is largely unknown how these anatomical and functional changes affect the integrity of cortical functional networks and how, in turn, changes in these networks affect visual functioning of patients. Understanding this may affect the applicability of future treatments; the modulation of brain activity could be integrated as an option for rehabilitation or training. Despite this, there have been very few studies assessing the local FC between specific regions of interest (ROI) in the brain of glaucoma participants. Moreover, in most studies, fMRI was applied while observers performed a task, which means that patients’ vision impairment could have affected both performance and imaging results (Gerente et al., 2015; Qing et al., 2010). It has been shown that using resting state (RS) blood oxygenation level-dependent (BOLD) signals is an effective way in assessing neuronal activity, and specifically, brain functional changes (van den Heuvel & Hulshoff Pol, 2010). RS is a particularly attractive approach to study populations with neuro-ophthalmic diseases like glaucoma. Since the patients do not need to perform any task, it can be used even when they are severely visually impaired as their impairment at the level of the eye will not affect the results (Carvalho et al., 2019; Frezzotti et al., 2014, 2016; Giorgio et al., 2018; J. Wang et al., 2017; J. Wang, Li, Wang, et al., 2016). By examining which areas of the brain are activated synchronously during RS scans, it is possible to cluster certain areas into resting-state networks (RSN) (van den Heuvel & Hulshoff Pol, 2010). These results indicate that the brain consists of multiple cooperative systems or networks. The known RSNs are named based on their function and include, amongst others the visual network (VN), the salience network (SN), dorsal attention network (DAN), and the default mode network (DMN). By examining FC within and between these predefined networks, it is possible to determine to what extent a pathology, such as glaucoma, affects the integrity of a participant’s brain networks.

The first aim of our study is to determine network integrity in glaucoma. For this, we characterized functional network activity over time and identified the most central brain areas (“hubs”) in glaucoma patients and control subjects. Fast eigenvector centrality mapping (fECM) was applied to each time course to attribute an eigenvector centrality (EC) value to each predefined region of interest (ROI). For each scan and for each group, the ROIs with EC values higher than the 95th percentile were considered the most central brain regions (“hubs”). Hubs for which we found a significant difference in EC between glaucoma and healthy in both scans were considered to provide evidence for network changes. Importantly, as mentioned before, by basing this analysis on resting-state (RS) fMRI data, we assess network integrity independently from the quality of the input coming via the eyes.

Our second aim is to investigate the different ways in which the VF could affect the hub function of networked brain areas. Therefore, in our study we defined three different behavioural scores based on VF sensitivity. These were calculated based on the notion that a brain region’s hub function might relate to: 1) the sensitivity of the worse eye – indicating disease severity, calculated by selecting the mean deviation (MD) value of the worse eye (WorseMD) 2) the sensitivity of both eyes combined – with one eye potentially compensating for loss in the other, calculated by taking the best value from overlapping locations in the total deviation maps of left and right eye and then calculating the mean of these values (Binocular Integrated Visual Field – BIVF); or 3) a difference in eye sensitivity – requiring additional network interactions, calculated as the absolute difference between the MD values of the right and left eye (AbsDiffMD). Next, we calculated the correlation between these behavioural scores and the centrality of the brain regions identified as being most influential in either glaucoma or control participants.

## 2. Material and methods

### 2.1 Study population

Twenty patients with primary open-angle glaucoma (POAG) and twenty-four control participants were included. Participants’ demographics are detailed in Table 1. Patients and controls did not differ regarding age (p=0.3). Participants were invited to the screening ophthalmic session only if they were MRI compatible: no past or current psychiatric disorder; no claustrophobia; no MRI incompatible implants; non-MRI safe tattoos; no use of recreational drugs or medications which may influence neurodegenerative progression. Prior to any examination, all participants signed an informed consent form. The study protocol was approved by the ethics board of the University Medical Center Groningen (UMCG). The study followed the tenets of the Declaration of Helsinki.

**Table 1 -.** Demographic and clinical information of participants. POAG: primary open-angle glaucoma; HC: healthy controls; VA: visual acuity; IOP: intraocular pressure (with treatment for glaucoma); OCT RNFL: Optical Coherence Tomography, retinal nerve fiber layer thickness; HFA MD: Humphrey Visual Field Analyzer, mean deviation; better/worse: eye selected based on better and worse HFA MD.

|  | **POAG (N=20)** |  | **HC (N=24)** |  |
| --- | --- | --- | --- | --- |
|  | Average | Standard Deviation | Average | Standard Deviation |
| Age (years) | 70.9 | 8.7 | 68.1 | 7.4 |
| Gender (M/F) | 10/10 |  | 16/8 |  |
| VA (R/L; decimal) | 0.9 / 0.9 | 0.4 / 0.3 | 1.1 / 1.0 | 0.2 / 0.3 |
| VA (better/worse; decimal) | 1.0 / 0.8 | 0.4 / 0.3 | 1.0 / 1.1 | 0.3 / 0.3 |
| IOP (R/L; mmHg) | 13.5 / 13.7 | 2.7 / 3.9 | 13.2 / 13.6 | 2.9 / 3.3 |
| OCT (R/L; μm) RNFL | 74.0 / 68.4 | 15.0 / 16.3 | 97.8 / 98.7 | 8.7 / 8.1 |
| OCT (better/worse; μm) RNFL | 74.3 / 68.1 | 15.6 / 15.6 | 98.6 / 97.6 | 9.1 / 9.5 |
| HFA MD (R/L; dB) | -7.0 / -9.0 | 8.7 / 8.3 | 0.0. / 0.1 | 1.3 / 1.0 |
| HFA ND (better/worse; dB) | -4.0 / -12.0 | 5.2 / 9.2 | 0.5 / -0.4 | 0.9 / 1.2 |

### 2.2 Ophthalmic Data Collection

All subjects participated in an ophthalmic session, which included visual acuity (VA) examination, intraocular pressure (IOP) measurement, optical coherence tomography (OCT), and VF assessment using standard automated perimetry (SAP).

Glaucoma participants were required to have been clinically diagnosed in at least on their eyes with POAG and have a SAP mean deviation (MD) of −2 dB or lower (more negative), measured with the Humphrey Visual Field Analyzer (HFA; Carl Zeiss Meditec, Dublin, CA, USA) SITA FAST strategy with either the 30-2 or 24-2 grid. Participants were included only if their VF loss was due strictly to glaucoma. They also had to have abnormal values in at least one eye for the thickness of the retinal nerve fiber layer (RNFL), as assessed by OCT (Canon HS-100, software version 4.1.0, Tokyo, Japan).

Healthy participants were required to have a best-corrected VA of at least 0.1 logMar (0.8 decimal) in both eyes. Their IOP had to be below 21 mmHg, as assessed with a non-contact tonometer (Tonoref II, Nidek, Aichi, Japan). VF integrity was checked using the HFA (Carl Zeiss Meditec, Dublin, CA, USA) SITA FAST strategy with either 30-2 or 24-2 grid, with the MD required to be higher than −2 dB. Their OCT assessment had to show a normal thickness of the RNFL in both eyes (no clock hours below 1st percentile allowed).

### 2.3 MRI and fMRI data acquisition

Scanning was carried out on a 3 Tesla Siemens Prisma MRI-scanner using a 64-channel receiving head coil. A T1-weighted scan (voxel size, 1mm^3^; matrix size, 256 x 256 x 256) covering the whole-brain was recorded to chart each participant’s cortical anatomy. Padding was used for a balance between comfort and reduction of head motion. The functional scans were collected using standard EPI sequence with 260 volumes (TR, 1350 ms; TE, 30 ms; voxel size, 3mm^3^, flip angle 68; matrix size, 94 x 94 x 45). Two fMRI resting-state scans were taken in a room in complete darkness for each participant. During both resting-state functional scans, participants were asked to close their eyes, relax, not think about anything in particular, and to try not to move.

### 2.4 fMRI data analyses

Image pre-processing, FC, ECM and statistical analyses were performed using SPM12 (Wellcome Department of Imaging Neuroscience, London, UK), fastECM toolbox (Wink et al., 2012), MarsBaR Region of interest (ROI) toolbox (http://marsbar.sourceforge.net/) and customised scripts, implemented in Matlab 2016b (The Mathworks Inc., Natick, Massachusetts). A related toolbox containing the code will be made available via the website www.visualneuroscience.nl.

#### 2.4.1 Image preprocessing

For each subject, the structural magnetic resonance image was co-registered and normalised against the Montreal Neurological Institute (MNI) template and segmented in order to obtain white matter (WM), grey matter (GM) and cerebrospinal fluid (CSF) probability maps in the MNI space. FMRI data were spatially realigned, co-registered to the MNI-152 EPI template and subsequently normalised utilising the segmentation option for EPI images in SPM. All normalised data were denoised using ICA-AROMA (Pruim et al., 2015). Additionally, spatial smoothing was applied (8 millimetres) to the fMRI data. No global signal regression was applied.

Based on the Power atlas coordinates (Power et al., 2011), 11 functional networks and associated 232 regions of interest (ROI) were defined (5 mm radius) using the MarsBar ROI toolbox for SPM12 (Brett et al., n.d.). For each ROI, a time-series was extracted by averaging across voxels per time point. Due to their related functions, some of the networks were excluded or modified: the sensory/somatomotor hand and mouth networks were combined into one network, the cerebellar network was excluded since our analysis focused more on cortical structures and the uncertain network was completely excluded from the analysis due to interpretation difficulties – due to the nature of uncertain networks, it is unknown what function each specific ROI pertains to. This left 131 ROIs spread over seven networks: DMN, memory network, visual network, salience network, ventral attention network and dorsal attention network.

#### 2.4.2 Prewhitening

To facilitate statistical inference, data were “pre-whitened” by removing the estimated autocorrelation structure in a two-step GLM procedure (Bright & Murphy, 2015; Monti, 2011). In the first step, the raw data were filtered against the 6 motion parameters (3 translations and 3 rotations). Using the resulting residuals, the autocorrelation structures present in the data were estimated using an Auto-Regressive model of order 1 (AR(1)) and then removed from the raw data. Next, the realignment parameters, white matter (WM) and cerebrospinal fluid (CSF) signals were removed as confounders on the whitened data.

#### 2.4.3 Functional connectivity analysis

For each participant, the pairwise temporal Pearson correlation between ROIs was calculated and a Fisher’s z-transformation was applied. The ROI’s z-values (hereafter: FC values) were averaged across participants. Then, the median group FC-values were used for the whole-brain analysis and used to compare the FC-values between groups using a family-wise error corrected (FWE) permutation test. Permuting the participant’s group labels was repeated 10000 times per participant, p≤ 0.05 was considered statistically significant.

#### 2.4.4 Eigenvector Centrality analysis

To determine the most important hubs of the predefined networks, fast ECM (Wink et al., 2012) was performed on the defined ROI time course data per subject. The ECM method builds on the concept of node centrality, which characterizes functional networks active over time and attributes a voxel-wise centrality value to each ROI. Such a value is strictly dependent on the sum of centrality properties of the direct neighbor nodes within a functional network. In the fast ECM toolbox (Wink et al., 2012), ECM is estimated from the adjacency matrix, which contains the pairwise correlation between the ROIs. To obtain a real-valued EC value, we added +1 to the values in the adjacency matrix. Several EC values can be attributed to a given node, but only the eigenvector with the highest eigenvalue (EV) will be used for further analyses. The highest EV values were averaged across subjects at group level. Based on these values, influential ROIs, i.e. the hubs (Betzel et al., 2016; Mišić et al., 2015; van den Heuvel & Sporns, 2011), can then be identified. Per group, only ROIs with EC coefficients higher than the 95th percentile (5% highest) were considered the most central and therefore used in the following analyses. Based on literature, the 95th percentile was chosen as an arbitrary threshold (Invernizzi et al., 2019). Furthermore, we quantify possible differences between influential hubs across groups by permuting labels with 10000 time repetitions. ROIs with p≤ 0.05 were considered statistically significant and therefore, categorized as well as influential hubs. FWE correction was applied for the number of group level comparisons, but not for the total number of ROIs analyzed.

In addition, a proxy distribution for the null hypothesis (H0) was obtained by generating 1000 times surrogate BOLD time series using the iterative amplitude adjusted Fourier transform method (iAAFT; (Räth & Monetti, 2009; Schreiber & Schmitz, 1996). In this way, correlations between ROIs were removed; the null distribution represents the amount of centrality obtained when no functional communication is present in the brain. Note that the null distribution of the ECM is not centered at zero, as EC values are forced to be positive real-valued. To define the confidence intervals of each EC value estimated per ROI, a bootstrap technique (across time-point) was used at group level in parallel to resample the filtered fMRI data. To support visualization, a Gaussian distribution was fitted to both bootstrap and surrogate distributions.

### 2.5 Behavioural scores

For all participants, the behavioural scores were derived from the HFA original output by: 1) selecting the mean deviation (MD) value of the worse eye (WorseMD; glaucoma: −12.11 [IQR 13.60] dB, controls: −0.31 [IQR 0.82] dB); 2) taking the highest value from overlapping locations in the total deviation maps of the left and the right eye and then calculating the mean of these values (Figure 1; Binocular Integrated Visual Field – BIVF; glaucoma: −2.63 [IQR 3.45] dB, controls: 0.79 [IQR 0.37] dB; (Crabb & Viswanathan, 2005); 3) calculating the absolute difference between the MD values of the right and left eye (AbsDiffMD; glaucoma: 4.82 [IQR 6.65] dB, controls: 0.75 [IQR 0.59] dB).

**Figure 1 -.**
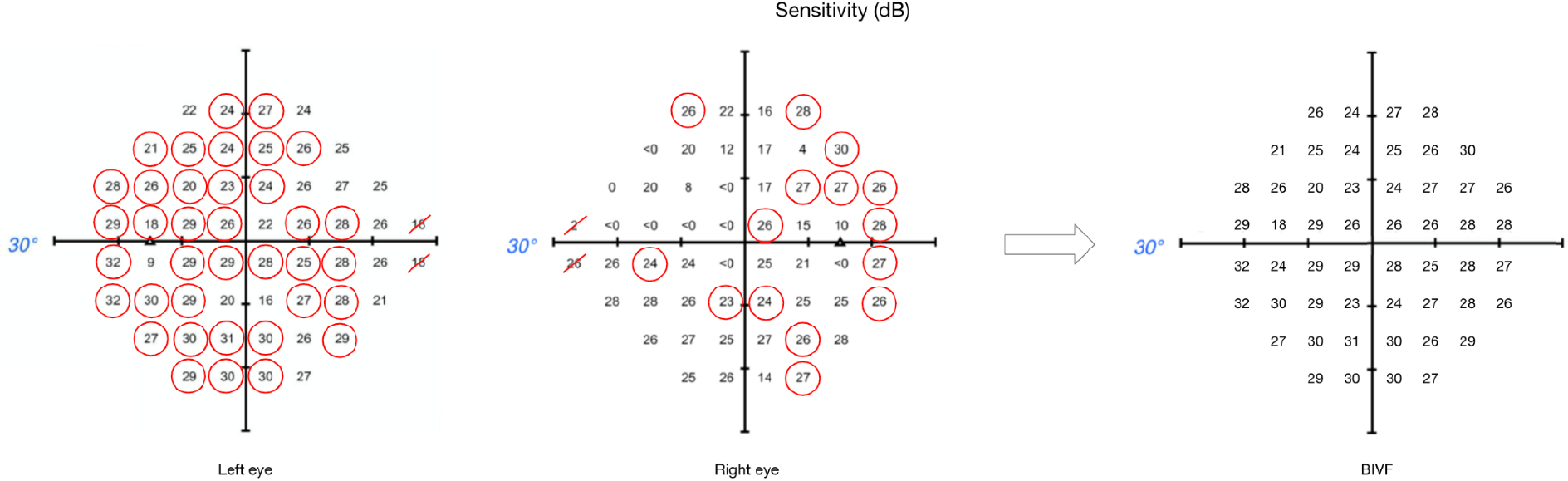
Binocular Integrated Visual Field calculation. This image shows the sensitivity maps from the original HFA output, for both left and right eyes respectively, and the resulting map for the BIVF. For each overlapping location between left and right eye, the highest value was selected (circled in red). The BIVF was calculated as the average of the selected values from the sensitivity map.

Finally, each derived behavioural score was correlated using Pearson’s correlation with EC value estimated per ROI. Only ROIs classified as influential hubs on both RS scans were considered in this analysis. Correlation with p≤ 0.05 was considered statistically significant.

## 3. Results

### 3.1 Whole Brain Connectivity Analysis

To compare whole brain connectivity between control and glaucoma participants, we plotted the averaged FC scores across all ROIs (Figure 2, Panel A) and the average EC values per ROI in both groups (Figure 2, Panel B and C, for controls and glaucoma participants, respectively). No significant differences were found with any of these analyses (RS1: HC versus POAG, p=0.59; RS2: HC versus POAG, p=0.089).

**Figure 2 -.**
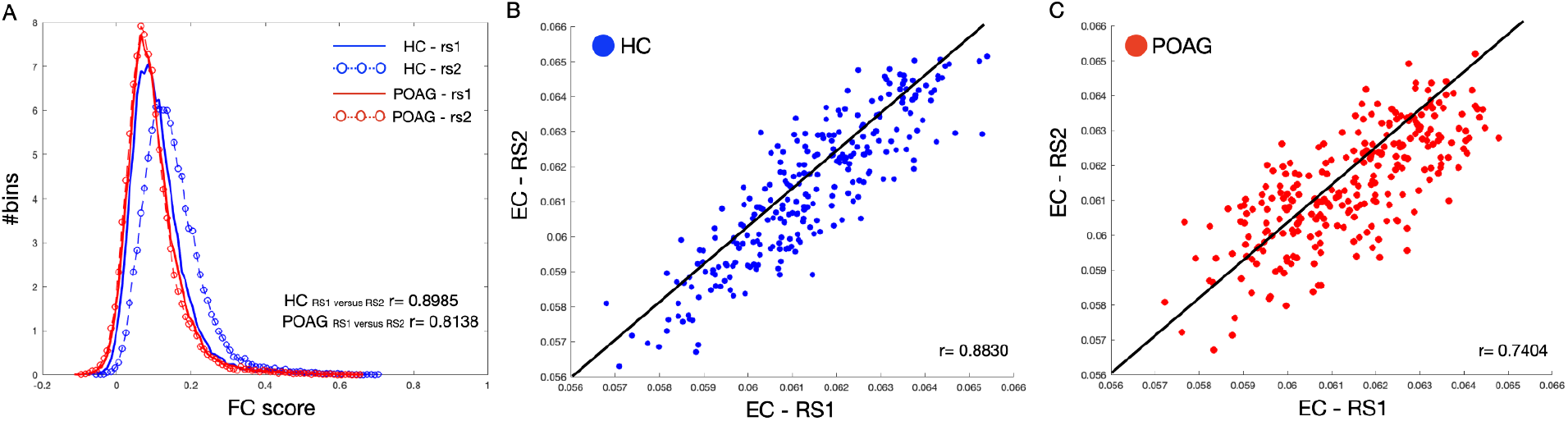
Test-retest evaluation between RS scans for both FC and EC values. Panel A shows the frequency distribution of the functional connectivity (FC) scores across all ROIs computed for both RS scans for control (blue line and circles) and POAG (red line and circles) participants. Panels B and C report the averaged EC values for control (blue) and POAG (red) groups, respectively. Each dot represents the average EC value for a single ROI defined based on the Power atlas. The black line represents the linear fit applied to the data.

Moreover, no significant differences were found with either the whole-brain-within or -between functional connectivity network analyses for controls and glaucoma groups (for further analyses see the supplementary Figures 1S, 2S and 3S).

### 3.2 Identification of influential brain areas

Based on the EC values averaged across participants, we identified the 5% most central ROIs per group, which we refer to as ‘hubs’ (Figure 3). For controls, hubs belonged to the Visual, Salience, Ventral and Dorsal Attention networks. For POAG, hubs belonged to the Visual, Salience, Default Mode, Ventral and Dorsal Attention networks. Based on the first RS scan, differences in EC between the POAG and controls did not reach significance for any of these hubs. In the second RS scan, differences reached significance for two hubs, both belonging to the Salience Network (Insula, p=0.023; Frontal Middle Gyrus, p=0.016 – Supplementary material, Figure S5).

**Figure 3 -.**
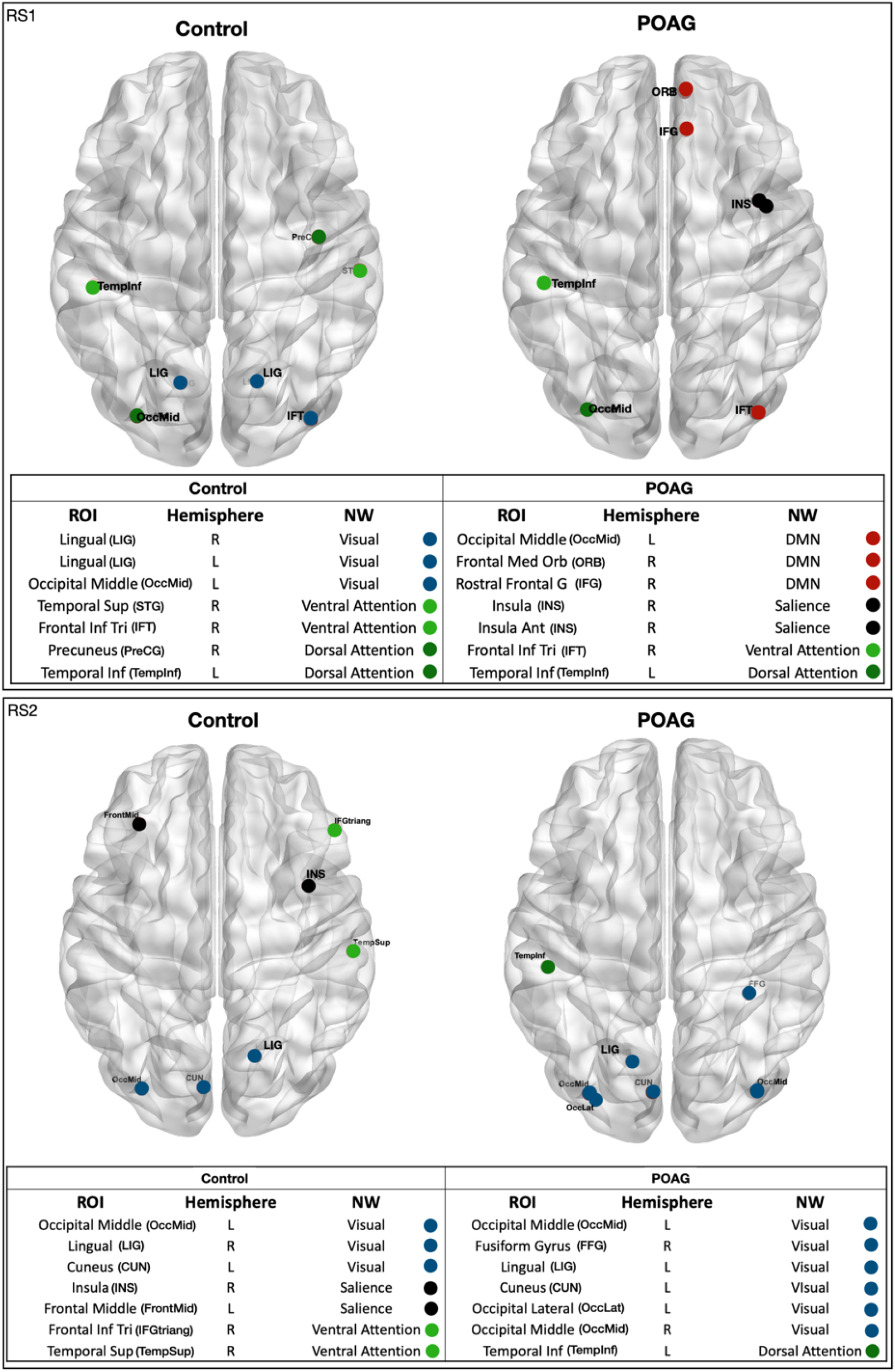
Functional network hubs identified in either of the two RS scans. The figure shows all identified hubs, i.e the ROIs with 5% highest eigenvector centrality in either of the RS scans in either group – healthy and POAG participants. The tables below the figures list the full name of the ROI, the ROI’s hemisphere, the name of the ROI’s functional NW and the NW color code.

To further explore and quantify the difference in EC value between the healthy and glaucoma participants, bootstrap and surrogate methods were applied to the EC values of the hubs identified in both RS scans. Figure 4 shows the bootstrapped and the surrogate EC distributions for a number of hubs present in both RS scans for healthy and glaucoma groups (Figure 4, RS1: Panel A; RS2, Panel B). Furthermore, we are reporting in supplementary material the EC distributions for the only two hubs that differences reached significance in RS2 (Insula and Frontal Middle Gyrus; Figure S5).

**Figure 4 -.**
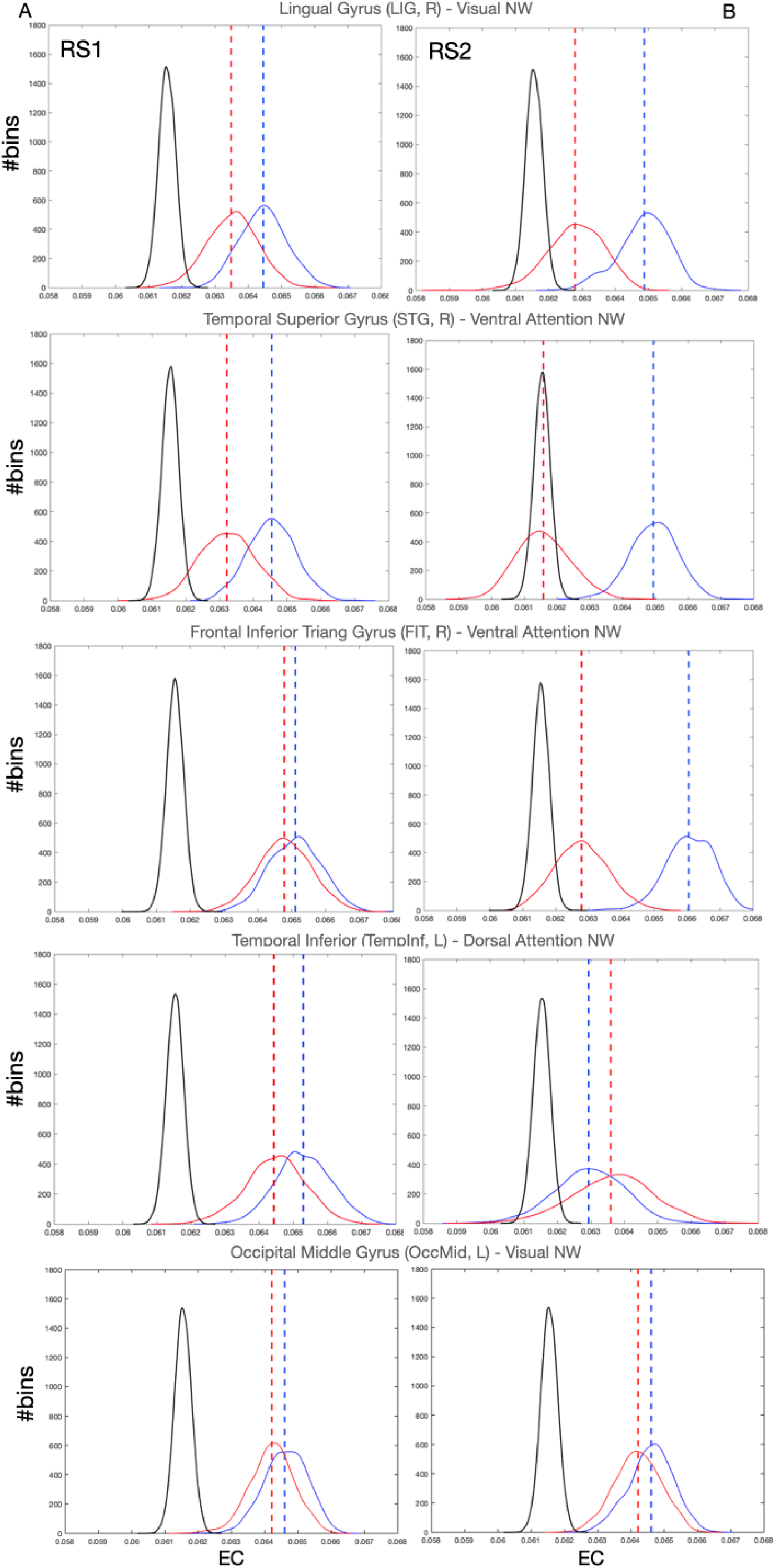
EC values of various hubs present in both RS scan of both POAG and controls groups. Eigenvector Centrality values of the hubs that were present in the RS scans for healthy and glaucoma groups (dashed lines), bootstrapped distributions (solid and dotted lines) and surrogate distributions for healthy and glaucoma (black lines) are reported for RS1 and RS2 in panels A and B, respectively. Note that the surrogate distributions for healthy and glaucoma participant groups are overlapping and indicated by H0 in the figure legend.

### 3.3 Correlations of EC of hubs with behavioural scores

For all identified hubs, based on the data of the first RS scan we found significant correlations between the EC values of the right Lingual Gyrus (LIG) and the BIVF (p=0.037), the AbsDiffMD (p=0.036), and the WorseMD (p=0.041) and (Fig. 5, left panels). Based on the data of the second RS scan, of these correlations, the one between the right LIG and the BIVF reached significance again (BIVF p=0.048; WorseMD p=0.228; AbsDiffMD p=0.690; Fig. 5, right panels). In other words, only the correlation between the right LIG and BIVF resulted in a reproducible effect. The right Lingual Gyrus belongs to the Visual Network (Power et al., 2011). In the supplementary materials, we report the bootstrapped and surrogate distributions for the hubs that showed no significant correlation between the EC values and the behavioural scores are shown (Figures S4).

**Figure 5 -.**
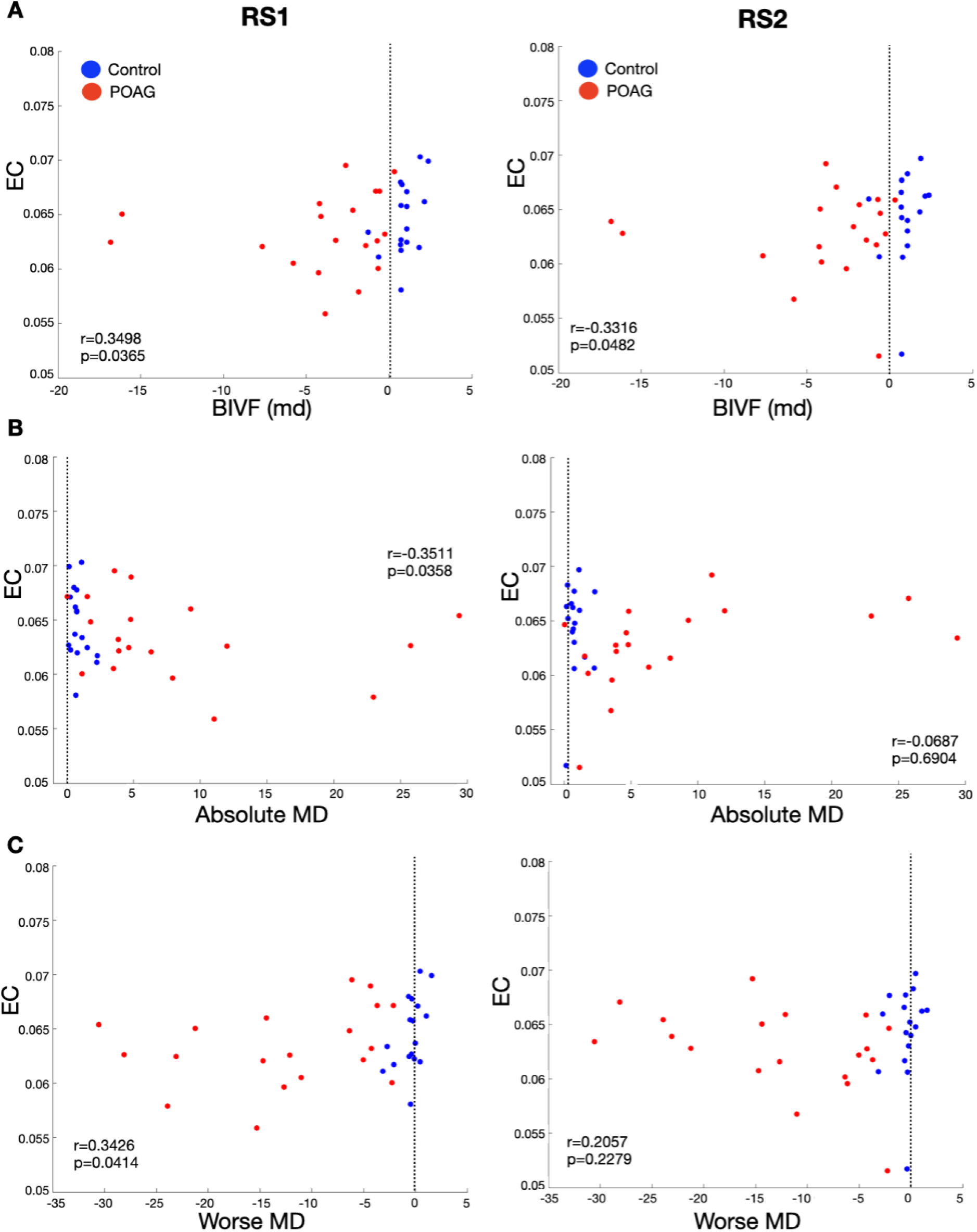
Correlation between the EC value of the Lingual Gyrus and obtained behavioural scores. Panel A,B and C report the Spearman’s correlation between the three behavioural scores and the EC values of the Lingual Gyrus estimated based on RS1 and RS2 scans. The black line indicates the zero for each behavioural score. For the behavioural scores: BIVF represents the integration for the MDs of left and right eye; Absolute Difference MD, the absolute difference between left and right eye MDs; Worse MD, which is the worse MD score between the left and right eye.

## 4. Discussion

This study has two main findings, one at the global and one at the local brain level. First, in glaucoma, global brain network communication is preserved. We conclude this based on an absence of whole brain differences in functional connectivity between glaucoma participants and controls. Secondly, in glaucoma, local brain hub connectivity relates to functional visual abilities. We conclude this because the hub function of the Lingual Gyrus is associated with the sensitivity of the integrated VF of both eyes. Below, we discuss these conclusions in more detail as well as the implications for understanding cortical functioning of glaucoma patients.

### Global brain network communication is preserved in glaucoma

No functional connectivity disruptions were found either within- or between-functional networks of glaucoma participants and controls. This is consistent with a previous study (J. Wang, Li, Wang, et al., 2016). Since global disruption of functional connectivity has been found in other syndromes such as Alzheimer’s disease (Pievani et al., 2011) and Parkinson’s disease (Pievani et al., 2011), these results support the idea that glaucoma has a different neuropathogenesis. Several other studies found functional changes at the level of single brain areas in glaucoma, however some of these studies investigated just a few specific vision-related brain areas, or focused only on FC within a specific set of networks, not between different networks (Dai et al., 2013; Frezzotti et al., 2014; Giorgio et al., 2018; Li et al., 2014; Qu et al., 2020; Song et al., 2014; Trivedi et al., 2019; J. Wang et al., 2017).

In our study, we found that, based on one of our RS scans, the centrality of the insula and the frontal middle gyrus, both part of the Salience Network, differed between glaucoma participants and controls. Yet, this finding did not replicate in the other scan. Overall, we conclude that the evidence for changed FC or centrality in glaucoma is weak at best.

### Reduced centrality of the LIG is associated with reduced functional vision

Based on a set criterion for their EC value, we identified a number of brain areas as being hubs. Of these hubs we found that the centrality of the LIG related consistently to the sensitivity of the VF of participants. While the EC of a few other hubs (Insula and Frontal Middle Gyrus) showed a correlation with one of the behavioural VF indices in either one of the scans, only the correlation of the EC of the LIG with the sensitivity of the binocular integrated VF (BIVF) reproduced in both scans. We can understand this result based on two previous findings. First, the LIG is known for its role in visuospatial processing and topographical recognition. It was found that lesions in this area affect the ability of patients to orient themselves (Mendez & Cherrier, 2003; Takahashi & Kawamura, 2002). Secondly, glaucoma patients often report issues in orientating and moving in their surroundings. This self-perceived visual disability was shown to relate most strongly to their peripheral vision loss when expressed in terms of the BIVF score (Crabb & Viswanathan, 2005). This implies the centrality of the LIG directly relates to this functional visual ability. It remains to be determined whether this relationship also implies local reorganization of the LIG. The reduced centrality of the LIG may either be a consequence of patients’ reduced visual input, mediated by their reduced orientation and mobility performance or a consequence of both. Irrespective, the fact that we consistently find this relationship based on RS data acquired in the absence of visual stimulation implies this altered centrality of the LIG has a permanent basis, implying brain reorganization has occurred.

### Clinical implications

The results in this study imply that network functionality poses no limit to future treatment options. The central role of the LIG suggests that future therapies, such as brain stimulation, could focus on this brain area, as its modulation may benefit glaucoma patients (Sabel et al., 2020). Moreover, the notion that orientation and mobility performance are also part of this equation could support the introduction of cognition-based mobility training in the treatment (Gunn et al., 2019; Virgili & Rubin, 2010).

### Limitations and future studies

In this study, we have grouped all glaucoma participants together, from early diagnosed ones with a very mild deficit to those with severe vision impairment. Consequently, in our glaucoma group, some participants had VFs that were mostly intact, while others had visual losses that caused a near complete blindness in the eye. While this was beneficial for our ability to relate centrality to functional vision, it makes it impossible to find more subtle patterns of reorganization. It is known that in early POAG, the brain goes through a stage where brain structures become larger before they atrophy (Williams et al., 2013), the result of this being a compensated change where neural activity is initially suppressed and then enhanced at a later stage in the disease (Johansen-Berg, 2007). Thus, it might be beneficial to further investigate how the different stages of glaucoma affect functional connectivity, again in association with patients’ behavioural scores. Finally, our present cross-sectional imaging approach does not allow us to establish causal relationships between the centrality of the LIG, the behavioural scores and the reduced mobility reported by patients. Future longitudinal studies in glaucoma (Haykal et al., 2021) could perhaps fill this void.

### Conclusion

We found no consistent alterations in the global or local functional networks of glaucoma participants. However, our study did show that the integrity of the hub function of the LIG relates to patients’ functional vision as expressed in their BIVF score. Future studies are required to determine whether deficits experienced by glaucoma patients in orientation and mobility are related to this observation, and if so, whether they could be either a consequence or a cause of changes in this brain area.

## Meeting presentation

none.

## Acknowledgements

The authors thank Prof. Dr. Nomdo Jansonius for his help in verifying the quality of the ophthalmic patient data.

## Supplementary Material

**Figure S1 -.**
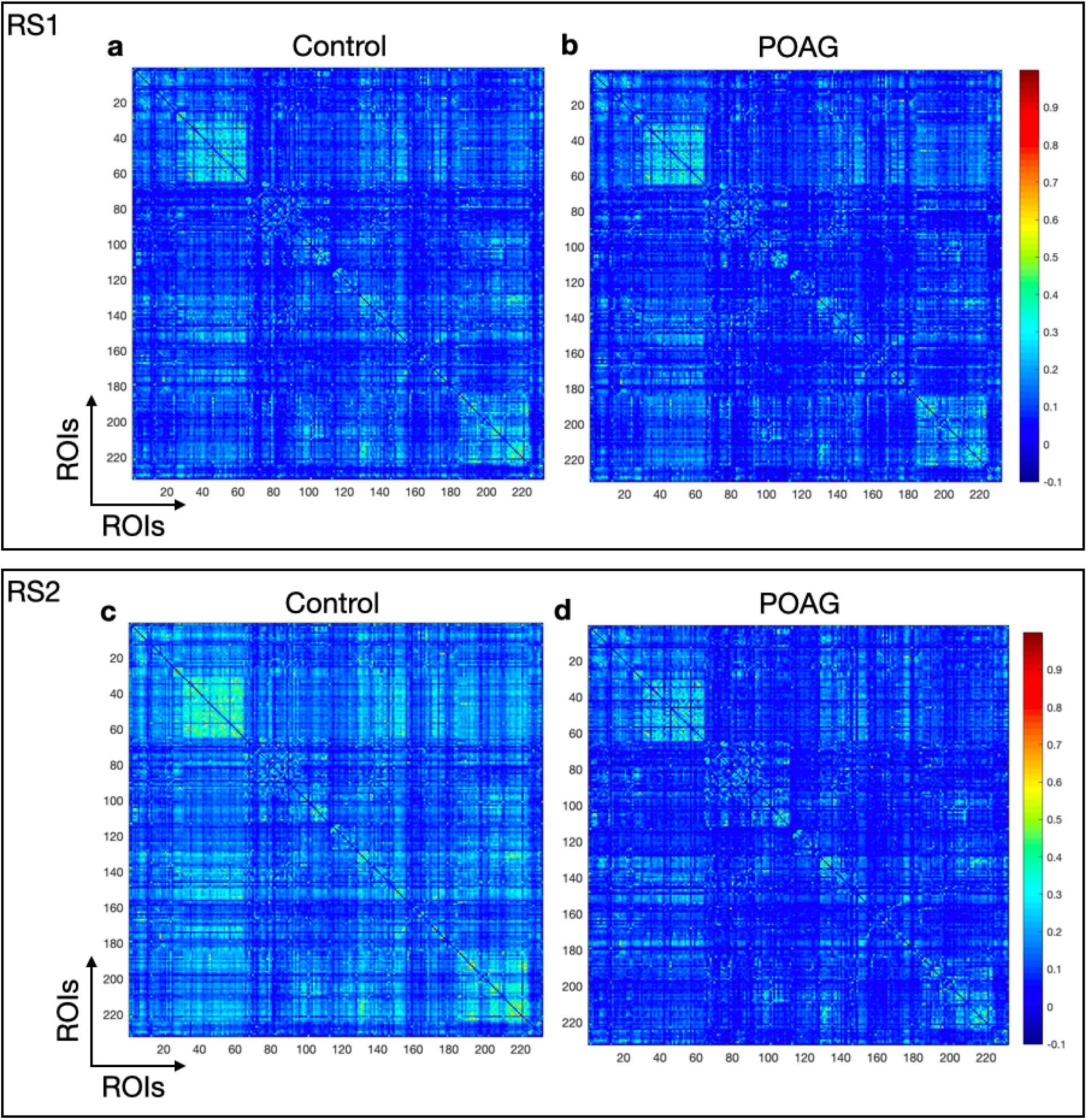
Whole brain functional connectivity matrices, based on the Power atlas. For each group, first all individual functional connectivity matrices were calculated by applying Fisher’s r-to-z transformation, which were then averaged across participants. Panels **a**, **b**, **c** and **d** show the mean inter-ROI correlation matrices for healthy and POAG participants, respectively.

**Figure S2 -.**
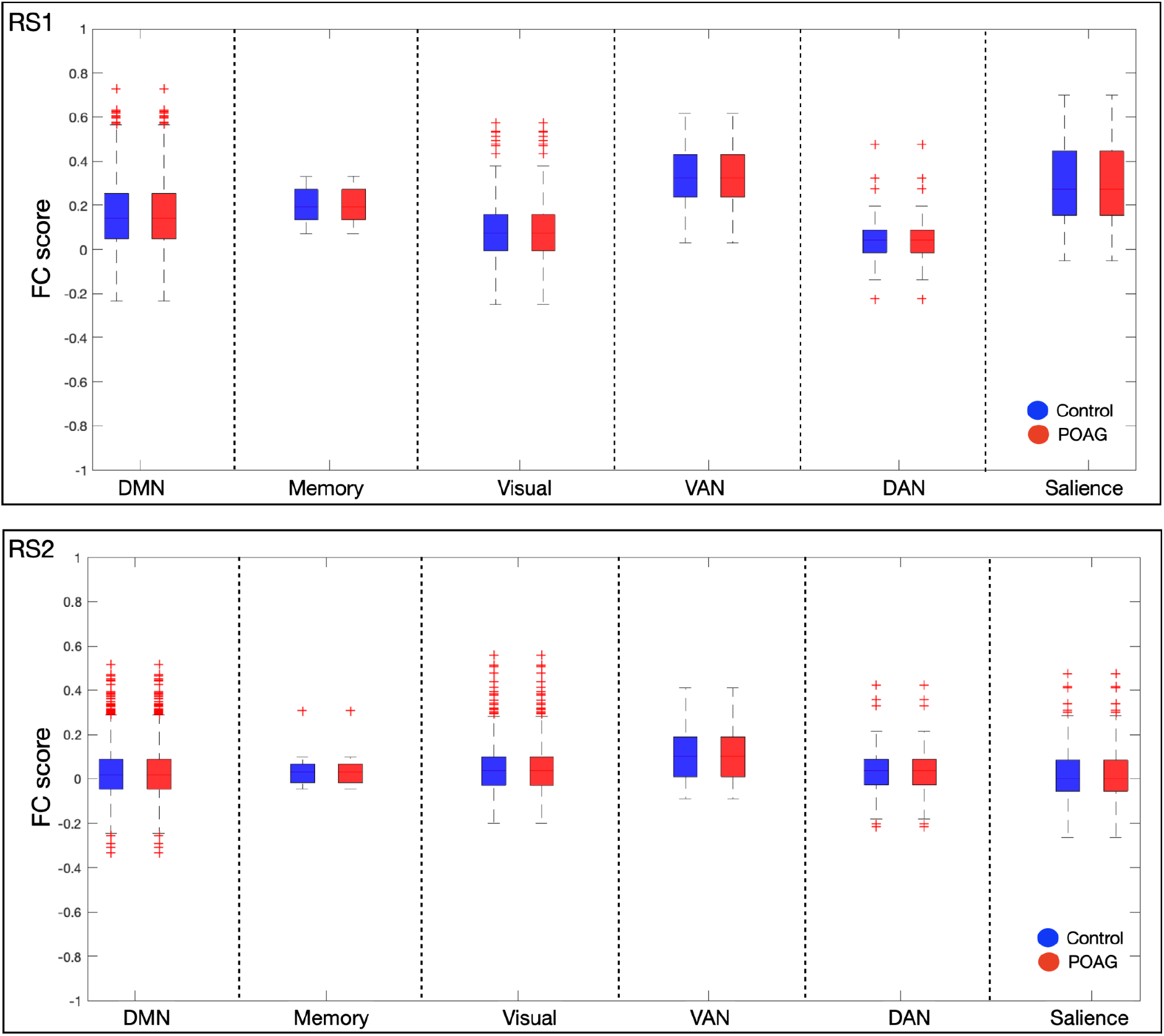
Intra-group brain functional connectivity analysis based on Power atlas. For each group, we averaged the functional connectivity score across all participants for the 6 predefined Functional networks. Top panel shows the intra-group analysis for RS1 while the bottom panel reports RS2

**Figure S3 -.**
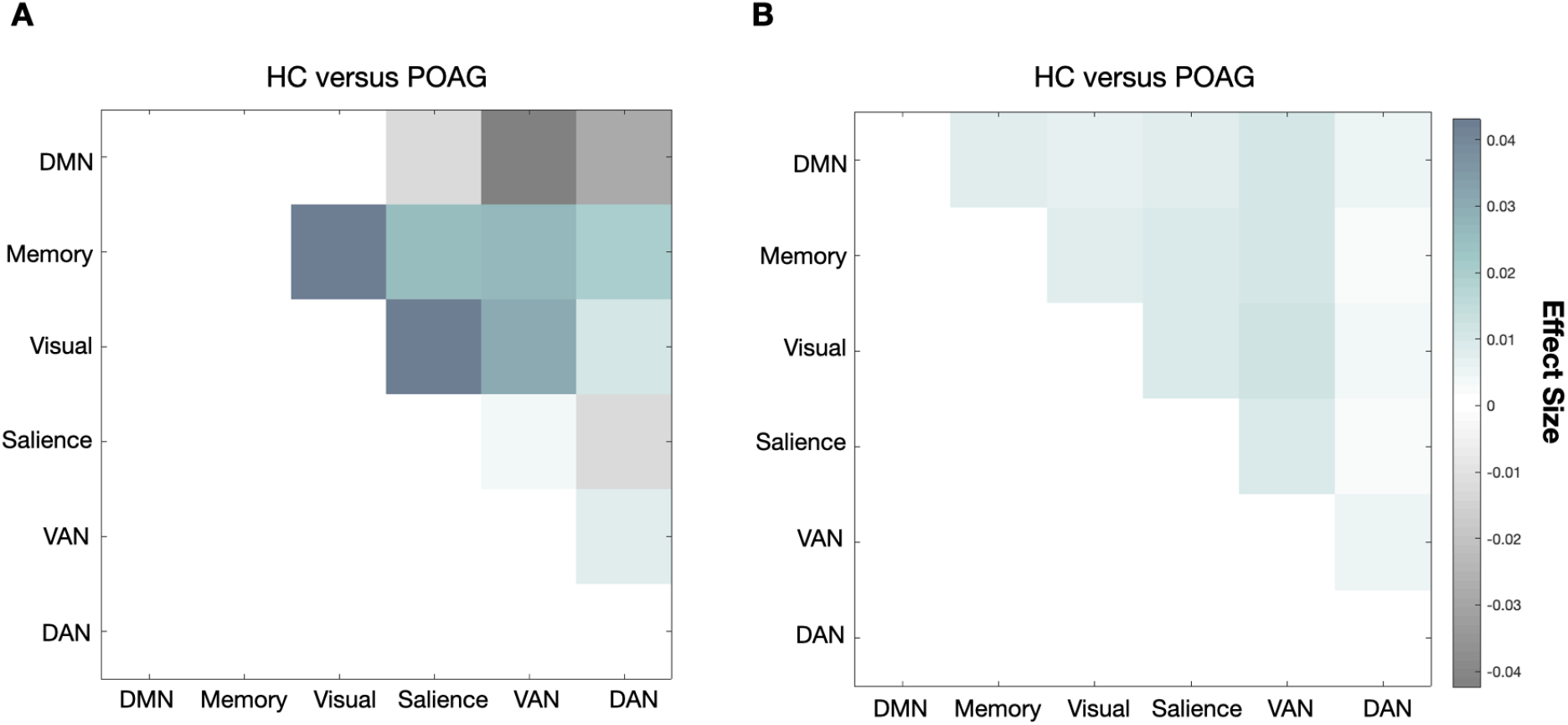
Functional connectivity effect size analysis between networks. For each group, we averaged the functional connectivity score across all participants and computed the effect size between healthy and POAG groups for the 6 predefined functional networks. Panels **a** and **b** report the effect size for RS1 and RS2, respectively.

**Figure S4 -.**
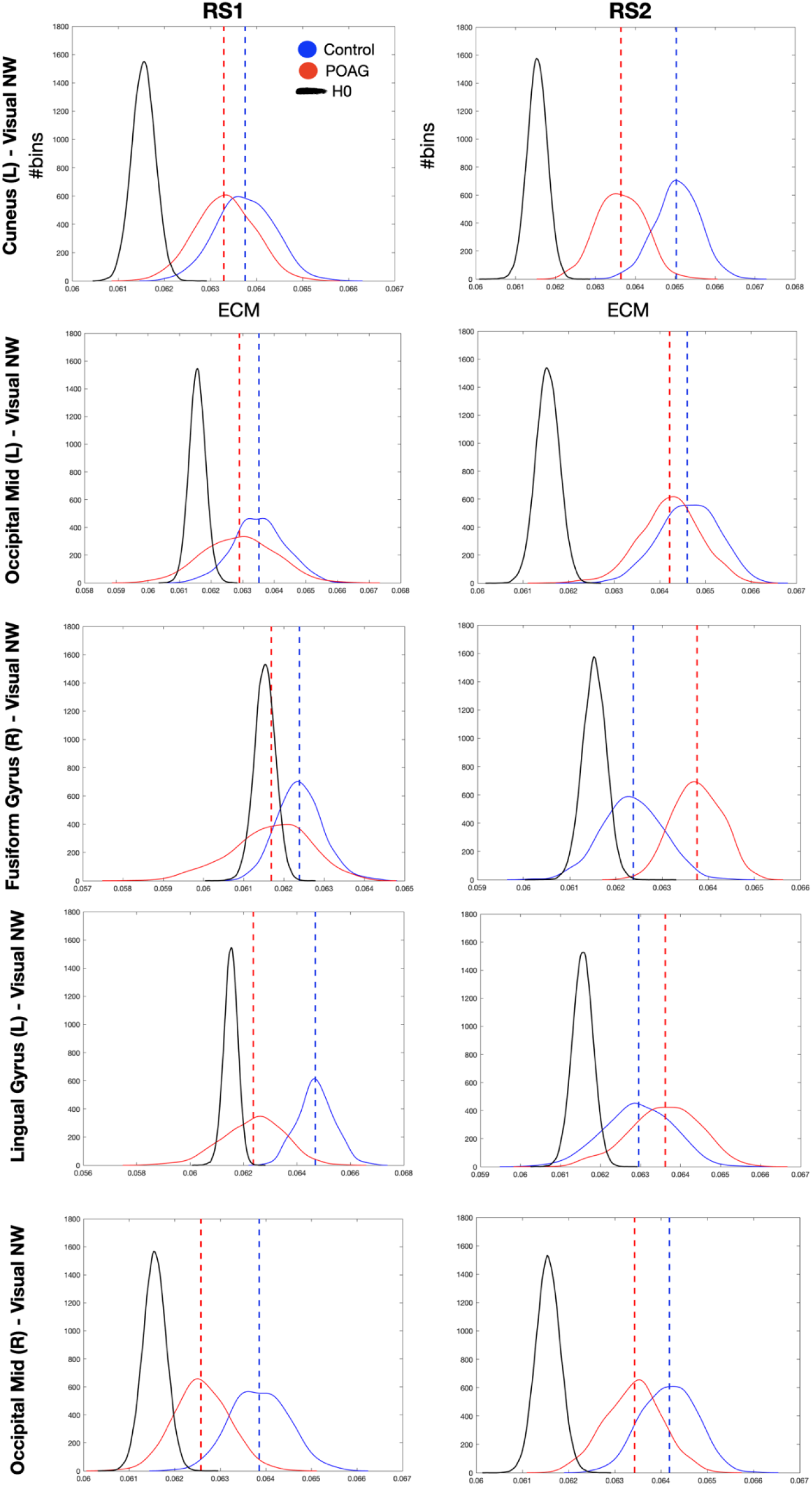
EC values computed on RS1 and RS2 across groups. Eigenvector values for healthy and glaucoma groups (dashed lines), bootstrapped distributions (solid and dotted lines) and surrogate distributions (back lines) are reported for the 5% most central hubs. Interestingly, these hubs did not significantly correlate with the behavioural scores even though they are part of the Visual Networks.

**Figure S5 -.**
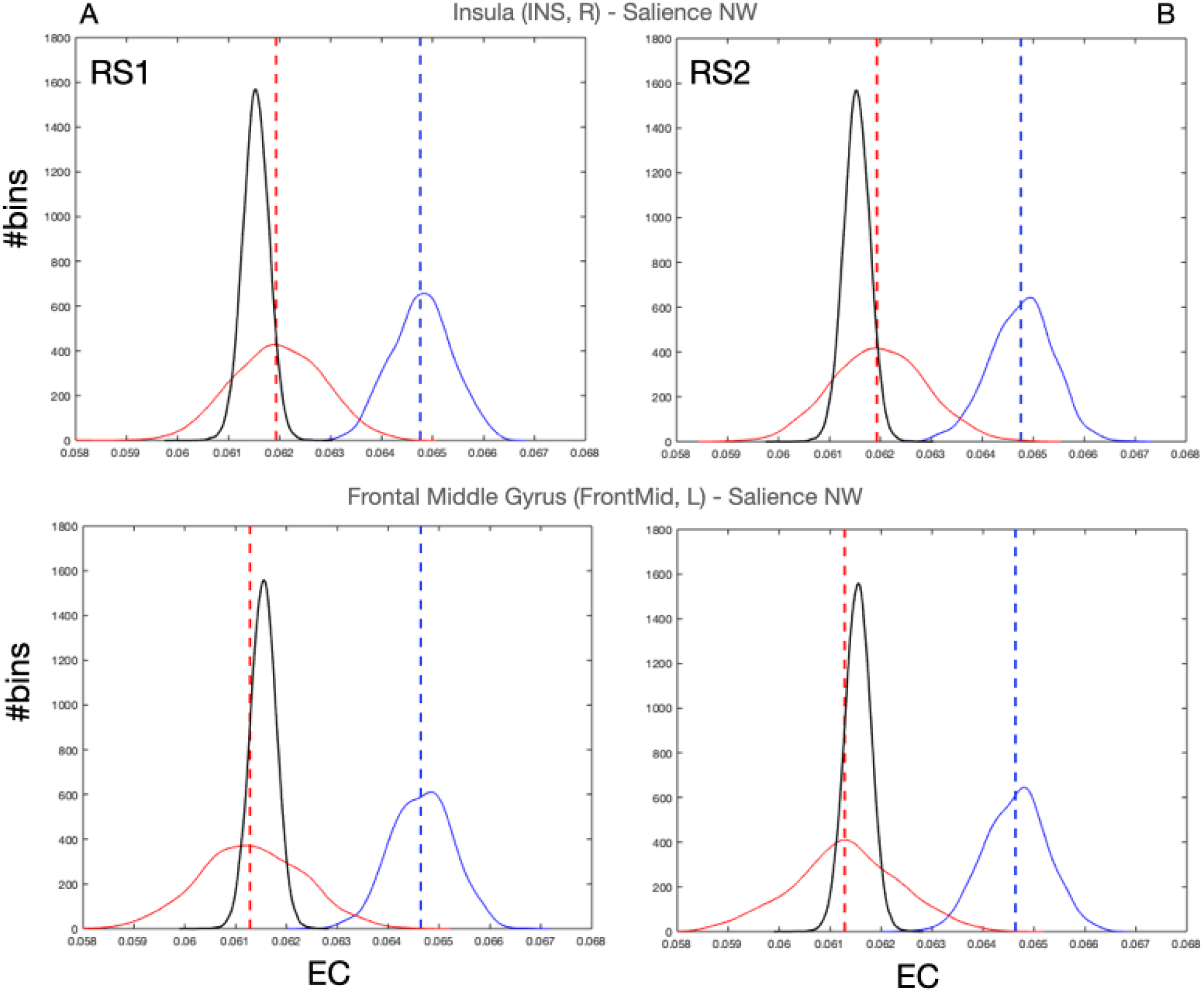
EC values of the only two significant hubs present in RS2 scan. Eigenvector Centrality values of the two hubs that reached significance in RS2 scan (dashed lines), bootstrapped distributions (solid and dotted lines) and surrogate distributions for healthy and glaucoma (black lines) are reported for RS1 and RS2 in panels A and B, respectively. Note that the surrogate distributions for healthy and glaucoma participant groups are overlapping and indicated by H0 in the figure legend.

